# Decoding Attended Spatial Location during Complex Scene Analysis with fNIRS

**DOI:** 10.1101/2022.09.06.506821

**Authors:** Matthew Ning, Meryem A. Yücel, Alexander Von Lühmann, David A. Boas, Kamal Sen

## Abstract

When analyzing complex scenes, humans often focus their attention on an object at a particular spatial location. The ability to decode the attended spatial location would facilitate brain computer interfaces for complex scene analysis. Here, we investigated functional near-infrared spectroscopy’s (fNIRS) capability to decode audio-visual spatial attention in the presence of competing stimuli from multiple locations. We targeted dorsal frontoparietal network including frontal eye field (FEF) and intra-parietal sulcus (IPS) as well as superior temporal gyrus/planum temporal (STG/PT). They all were shown in previous functional magnetic resonance imaging (fMRI) studies to be activated by auditory, visual, or audio-visual spatial tasks. We found that fNIRS provides robust decoding of attended spatial locations for most participants and correlates with behavioral performance. Moreover, we found that FEF makes a large contribution to decoding performance. Surprisingly, the performance was significantly above chance level 1s after cue onset, which is well before the peak of the fNIRS response. Our results demonstrate that fNIRS is a promising platform for a compact, wearable technology that could be applied to decode attended spatial location and reveal contributions of specific brain regions during complex scene analysis.

## Introduction

The human brain is an astonishingly powerful computational device, capable of feats yet to be matched by machines. One impressive example is the brain’s ability to selectively attend to specific objects in a complex scene with multiple objects. For example, at a crowded cocktail party, we can *look* at a friend and *hear* what they are saying in the midst of other speakers, music and background noise. Such multisensory filtering allows us to select and process important objects in a complex scene, a process known as Complex Scene Analysis (CSA) [1,2,3]. In stark contrast, millions of humans worldwide with a varying degrees of hearing losses and disorders, e.g., ADHD [4,5], autism [6,7], hearing losses [8], find such complex scenes confusing, overwhelming and debilitating. Thus, brain-computer interfaces (BCIs) and assistive devices for CSA have the potential to improve the quality of life for many humans.

CSA can be broken down into several distinct components: determining the spatial location of a target stimulus, segregating the target stimulus from the competing stimuli in the scene, and reconstructing the target stimulus from the mixture forming a perceptual object [9]. Recently, we proposed a brain inspired algorithm for auditory scene analysis based on a model of cortical neurons [10-12]. This algorithm has the potential to be applied in assistive devices for CSA. However, a critical piece of information required by the algorithm is the spatial location of the target stimulus. Thus, a portable and non-invasive technology that can decode the spatial location of the attended target stimulus during CSA would greatly facilitate the development of BCIs and assistive devices for CSA. In addition, a technology that provides insights into specific brain regions which play a significant role in decoding the attended location has the potential for advancing our understanding of fundamental brain mechanisms underlying CSA in both normal and impaired humans.

Previous studies have employed electroencephalography (EEG) to decode spatial location of auditory attention (for a review, see [13]). EEG is a portable, non-invasive technology that is potentially well suited for decoding. One important class of auditory attention decoding (AAD) algorithms is the stimulus-reconstruction method, which uses the envelopes of EEG signals to reconstruct the attended speech envelope. Then, the correlation coefficients of the reconstructed speech envelope with the speech envelopes of different speakers are calculated, and the highest one determines the spatial location [14-21]. Beside stimulus-reconstruction approach, there are AAD algorithms based on EEG signals themselves, including common spatial pattern [22], and convolutional neural network [23-25]. However, a limitation of EEG based approaches is that its poor spatial resolution makes it difficult to gain insight into specific underlying brain regions.

In contrast, fMRI has provided insights into brain regions involved in auditory-spatial short-term memory [26]. Specifically, auditory short-term memory tasks elicited bilateral hemodynamic responses in the transverse gyrus intersecting precentral sulcus [26,27]. fMRI also showed that with demanding auditory spatial short-term memory tasks visuospatial regions, e.g., intraparietal sulcus 2-4 and superior parietal lobule 1, can also be activated. In addition, non-visuospatial regions near these regions, e.g., anterior intraparietal sulcus (antIPS), lateral intraparietal sulcus (latIPS), and medial superior parietal areas, were also activated by the same tasks [26,27]. In separate studies, fMRI also showed STG/PT to be activated in sound-localization tasks [28,29]. In addition, dorsal frontoparietal region, including frontal eye field in the frontal cortex and posterior parietal cortex and intraparietal sulcus, are shown to be activated by both audio-spatial and visuospatial tasks in fMRI studies [30-32] and simultaneous magnetoencephalography (MEG)/EEG study [33]. Although fMRI has revealed a wealth of information regarding brain regions activated during a wide variety of tasks, one limitation of fMRI is that it not portable, making it difficult to apply in BCIs and assistive devices.

Both visual and auditory cues play an important role in CSA. One important observation made in the last few decades is that visual and auditory modalities can influence each other as shown in psychophysical, neurophysiological and neuroimaging studies. These studies suggest the idea that perception combines features from both visual stimuli and auditory stimuli to form a single auditory-visual object [9]. Both auditory and visual features can contribute cross-modally to enhance auditory or visual object formation, which in turn, can enhance attention operating on auditory [30,34-36] or visual objects [37,38]. Cross-modal influences have been demonstrated via activation or modulation in primary visual cortex by auditory stimuli [39], single-unit and local field potential modulation in primary auditory cortex by visual stimuli [40,41] and fMRI study on humans [42-45]. Thus, employing auditory-visual objects to investigate CSA has the potential to reveal additional aspects of CSA beyond paradigms that utilize auditory stimuli alone, e.g., EEG-based AAD algorithms.

An alternative technology that is both portable and has good spatial resolution is fNIRS, making it well suited for applications in BCIs and assistive devices, as well as revealing specific brain regions activated during tasks. Previously, fNIRS has been applied to classify different sound categories [46], to identify spatial locations of noise stimuli [47], characterize hemodynamic responses to varying auditory stimuli [48-50] and investigate informational masking [51,52]. However, to date fNIRS has not been applied to decode auditory-visual spatial attention during CSA. Here we investigate the capability of fNIRS in such a decoding task. We find that fNIRS provides a robust platform for decoding attended spatial location. Surprisingly, improvement in decoding performance is faster than anticipated given the time-scales of the underlying fNIRS signals. Finally, we also find that FEF is a critical brain region for decoding attended spatial location during CSA.

## Materials & Methods

This study employs continuous wave fNIRS (CW6, Techen System) using 690nm and 830nm wavelengths, with 50 Hz sampling rate. We used a size 56cm Landmark Cap (EasyCap, Herrsching, Germany) for all subjects. By fitting the cap on a head model marked with EEG 10-20 reference points, we positioned and secured rubber grommets with panel holes to the cap to hold both source and detector optodes. Multiple channels are supported with frequency multiplexing. The fNIRS recording software is synched to the stimulus presentation with Neurospec MMB70-817 triggerbox (Neurospec AG, Switzerland).

### Dataset

The videos used in the experiment are from AVSpeech, a publicly available dataset (Ephrat et al. 2018, https://looking-to-listen.github.io/avspeech/index.html). For target stimuli, each subject was shown 6 different speakers, each speaker having 5 different clips, all showing the speakers’ faces clearly. Each clip was shown at all spatial locations at equal frequencies. For masker stimuli, each subject was shown 12 different speakers which are separated from the pool of speakers designated for target stimuli. These 12 speakers also have 5 different clips. For both pools of targets and maskers, all are English speakers. Each subject was presented with different pools of speakers and maskers. For each video clip, the audio clip was extracted and passed through three different head-related transfer functions (HRTF) to create auditory virtualization, one for each location (center, 45° to the left, and 45° to the right). The HRTF filters are based on measurements using the KEMAR head model and are publicly available on https://sound.media.mit.edu/resources/KEMAR.html. For each location, the HRTF-filtered audio clip replaced the original audio waveform in the video clip. Thus, there is a total of 90 trials, 30 for each location. Stimuli delivery software was written with Psychtoolbox-3 library.

### Participants

12 adults with normal hearing (age 19-48) were recruited for this study in accordance with the Institutional Review Board of Boston University. A COVID-19 protocol was developed and strictly adhered to. Participants were screened to exclude those with neurological disorders. Participants were briefed and consented before partaking in this study and were compensated for their time.

### Experiment

A computer with three monitors was used for the experiment. The monitors were located at the three locations: center, 45° to the left, and 45° to the right (Fig. 1A). They all are equidistant from the center where the subject is sitting upright. The auditory stimuli were delivered via earphone (ER-1 Etymotic Research Inc.) with eartips (E-A-RLink 3A Insert Eartips). This study adopted event-related design. For each trial, a 2 seconds long audio-visual cue was delivered in the form of a white cross against a black background and a 2 kHz pure tone linearly ramped in the first 0.5 s. This audio-visual cue appeared randomly at one of the 3 spatial locations, indicating the location of the target speaker. The cue was followed by 3 videos, one for each location, one of which is the target speaker and the 2 remaining are the maskers. It’s followed by two multiple-choice questions, each question containing 5 possible choices. These multiple-choice questions are always displayed at the center location. The two questions are related to face identification and words identification, presented in that order. In the face identification task, the subjects are presented five different faces and is tasked with correctly identifying the face of the target speaker shown in the video. In the word identification task, the subjects are presented five different transcripts and is tasked with correctly identifying the words spoken by the target speaker in the video. Upon the completion of two questions, a blank black screen of jittered duration with uniform distribution between 14 and 16 seconds appeared. This is to increase the statistical power of GLM fitting by preventing fNIRS signals from synching with the timing of the trial. Thereafter, an instruction to press the space bar to begin the next trial is displayed at the center location. While the audio-visual cue and the video clip last 2 and 3 seconds respectively, the subjects have 20 seconds in total to answer both multiple-choice questions but can move on to the space bar instruction immediately after the questions are done. Lastly, the subjects are provided with chin rest in order to discourage head movements.

**Figure 1.**
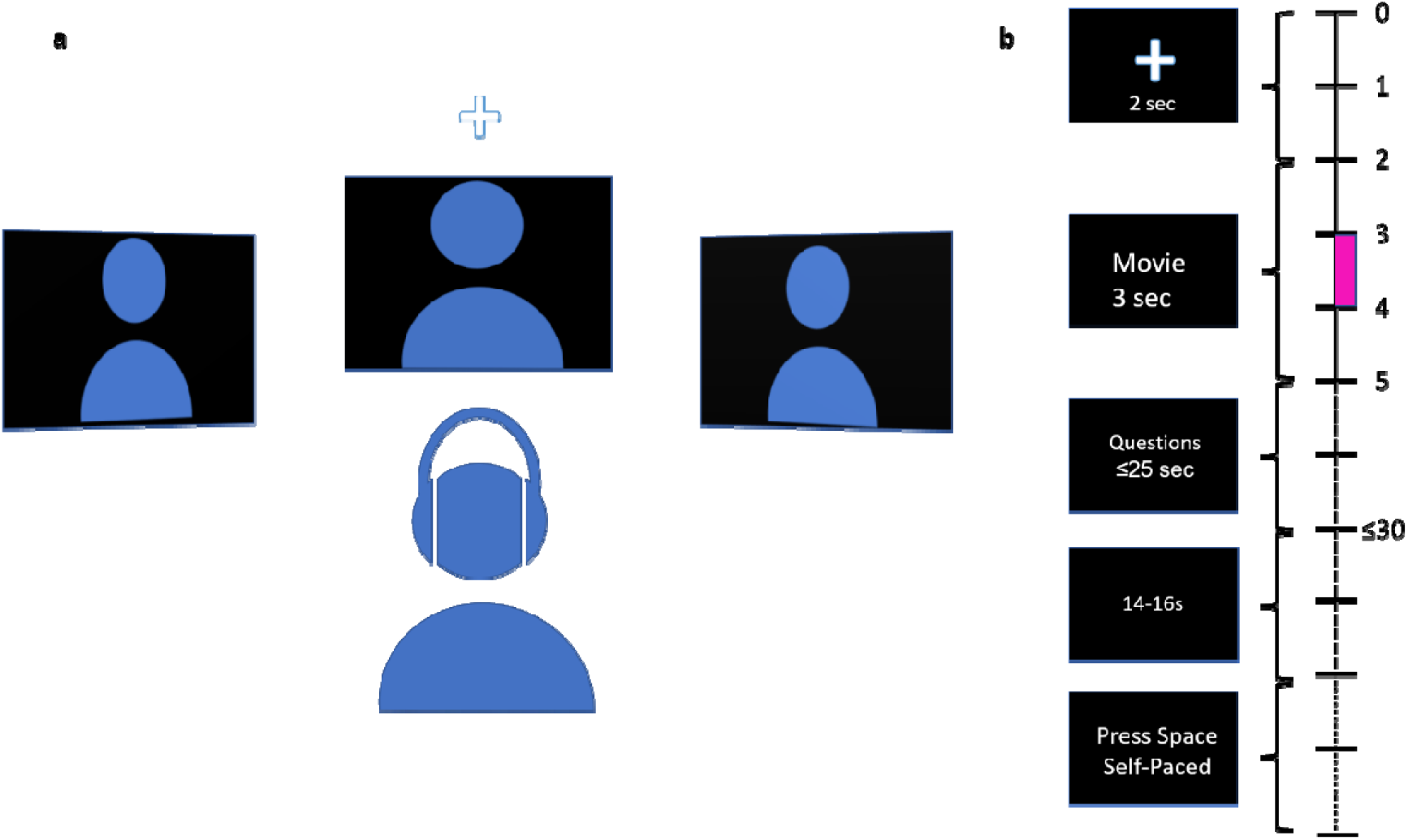
Schematic diagram of the experiment and trial. (a) Schematic diagram of experimental setup. Subject is seated in front of 3 spatially separated monitors, one at −45° to the left, one at 45° to the right, and one at 0° at the center. Audio is delivered via earphone. (b) Timeline of trial. The time at the right indicates the length of different parts of trial. A cue in the form of a white cross accompanied by a pure ramping tone of 2 kHz randomly appears at one of the three locations to indicate the location of the target stimuli at the start of the trial. Next, 3 video clips are displayed simultaneously for 3 seconds. Next, two multiple-choice questions are displayed at the center monitor, to be answered with keyboard. 1^st^ question is to identify the face of the speaker, 2^nd^ question is to identify the words spoken by the target speaker. Subject has up to 25 seconds to answer both questions. Upon the completion of the questions, it’s immediately followed by a blank screen of jittered duration between 14-16 seconds. Next, instruction to press the space bar is displayed at the center monitor to begin the next trial. Pink box in 3 ≤t≤4 is the decision window used in Figure 5, 6 and 7.

### fNIRS probe design

The probe was designed in AtlasViewer (https://github.com/BUNPC/AtlasViewer), an open-sourced MATLAB software and is shown in Figure 2. The probe contains 12 sources and 17 regular-separation detectors and 6 short-separation detectors, for a total of 30 regular channels (30 mm) and 6 short-separation (SS) channels (8 mm). The probe covers the following regions: intraparietal sulcus (IPS) 2, 3 and 4, superior precentral sulcus (sPCS), inferior precentral sulcus (iPCS), dorsolateral prefrontal cortex (dlPFC), frontal eye field (FEF), posterior superior temporal gyrus/planum temporale (STG/PT), anterior intraparietal sulcus (antIPS), superior parietal lobule (SPL1) including medial superior parietal lobule (mSPL), and transverse gyrus intersecting precentral sulcus (tgPSC). The probe was designed with multiple channels to cover variation in reported MNI coordinates of FEF.

**Figure 2.**
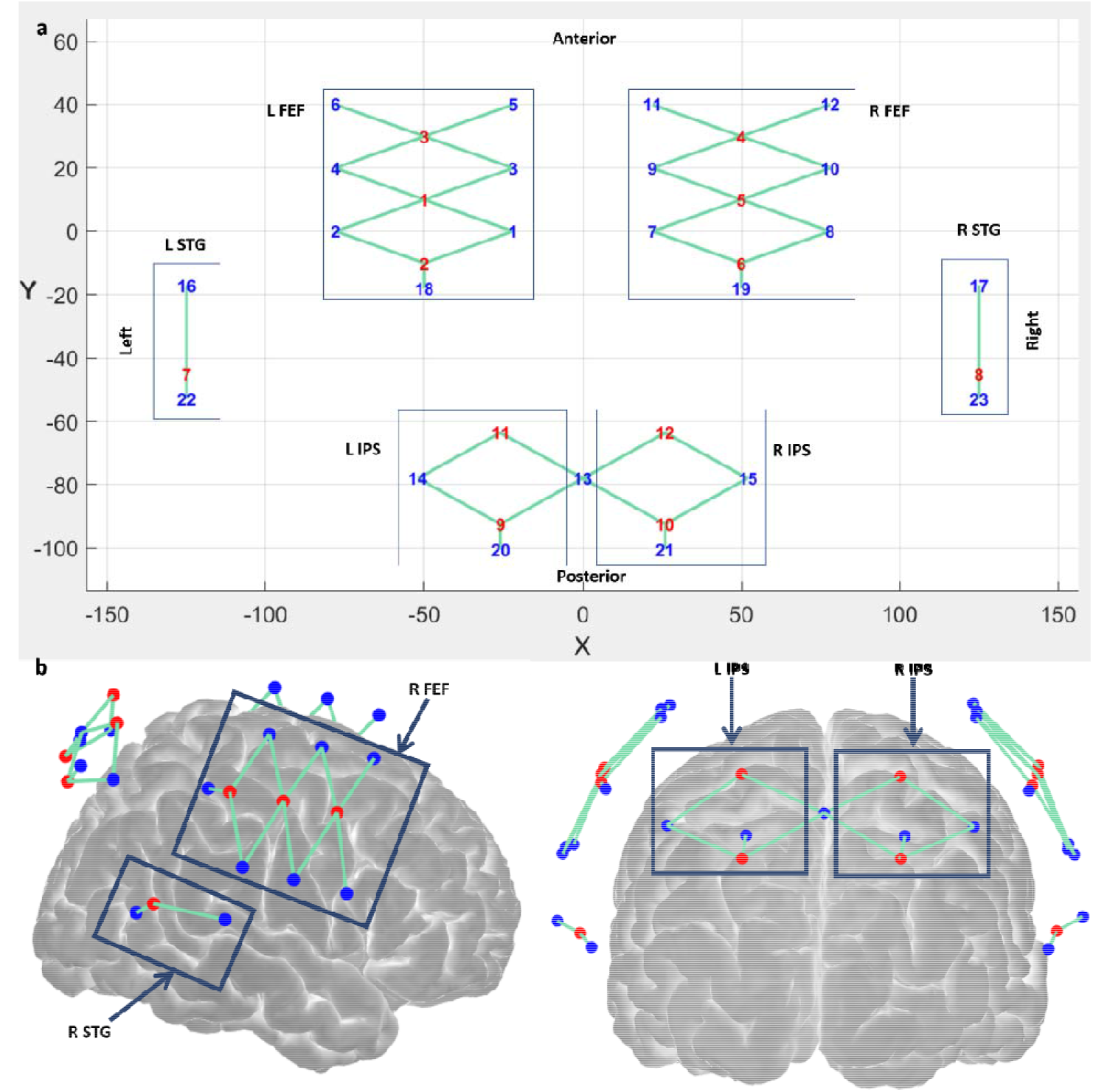
Probe Design. (a) 2D layout of probe design. The probe is subdivided into 5 regions: left frontal eye field (L FEF), right frontal eye field (R FEF), left posterior superior temporal gyrus (L STG), right posterior superior temporal gyrus (R STG), left intraparietal sulcus (L IPS) and right intraparietal sulcus (R IPS). Top side of the probe corresponds to the anterior side of the head, bottom is posterior. Red numbers represent source optoces, blue numbers represent detector optodes (b) 3D layout of probe design. Red circles represent source optodes, blue circles represent detector optodes, and turquoise lines represent channels.

### SNR

We excluded subjects that had at least 20 channels pruned using a cutoff of SNR = 1.5, where SNR is estimated as the mean divided by the standard deviation of the fNIRS signal.

### Grouping by Task Performance

We divided subjects into 2 groups, the high-performing group consisted of subjects that scored at least 80% correct on trials, and the low-performing group consisted of subjects that scored less than 80%. A trial is counted as correct if the participant correctly identified both the faces of speakers and the words spoken by the targeted speakers.

### fNIRS processing

For Fig. 3-7, raw light intensities were converted to optical densities using the mean of the signal as the arbitrary reference point. Next, motion artifacts in optical density (OD) were identified and corrected with targeted PCA before applying criteria for automatic rejection of stimulus trials [53]. Next, OD signals were band-pass filtered between 0.01 Hz and 0.5 Hz with a 3^rd^ order zero-phase Butterworth filter. Next, the filtered OD were converted to chromophore concentration changes. The differential path length factor is held fixed at 1 and the concentration unit is Molar*mm [54]. Finally, systemic physiological signal clutter was regressed out using a GLM with short-separation channels modeling the signal clutter. Each regular channel was assigned a SS channel with the highest correlation [55]. All were done using the Homer3 open source fNIRS analysis package and custom MATLAB scripts.

**Figure 3.**
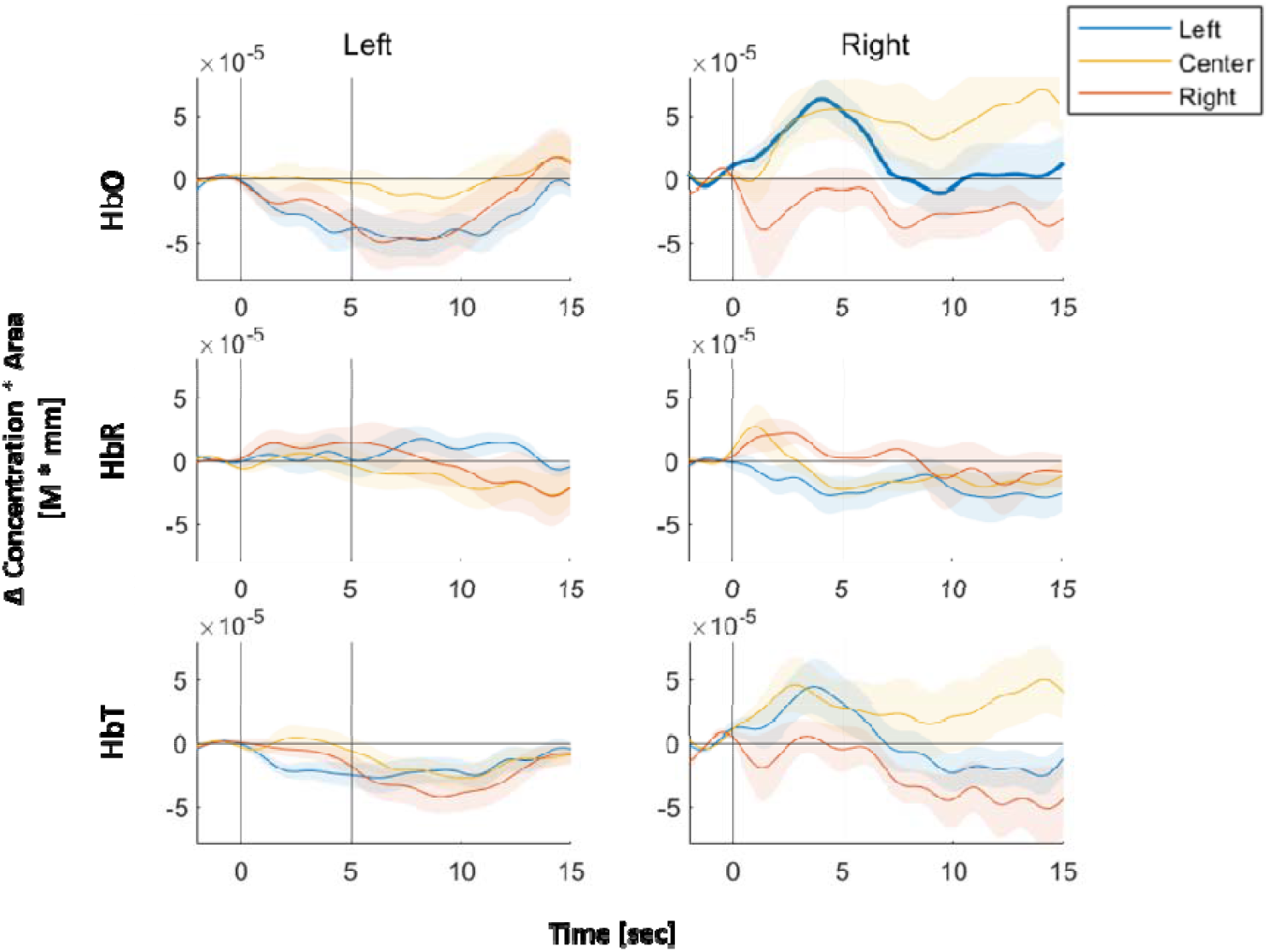
Hemodynamic Response Functions (HRFs) of FEF. Top panel represents Δ[HbO], middle panel represents Δ[HbR] and bottom panel represents Δ[HbT]. Left column represents channel covering FEF in the left hemisphere. Similar for right column. Blue line represents GLM-fitted HRF responses to left stimuli, red line for right stimuli, and yellow line for center stimuli. Shaded regions represent 95% confidence interval. Bold line indicates statistical significance in activation using one-sample two-tailed t-test. * indicates statistical significance in difference between left and center condition using two-sample two-tailed t-test, + for left vs right condition, and x for right vs center condition. In this case, there is no statistical difference between any pairs of spatial locations. The horizontal black line is the axis at 0 M*mm. Right FEF channel showed the highest correlation coefficient (R = 0.38, n = 9) between task performance and single-channel CV accuracies

For Fig. 8, in order to accurately assess the latency of the classification accuracy, the OD signals were band-pass filtered between 0.01 Hz and 10 Hz with a 3^rd^ order zero-phase Butterworth filter.

### GLM

For HRF modeling, we fitted the GLM model to each subject. The GLM fitting here uses two classes of regressors: HRF for each condition and the systemic signal clutter. The temporal basis functions used to model the HRF is a sequence of 16 Gaussian functions 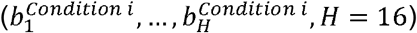, spaced 1 second apart (Δ*t* = 1*s*) and each with a width of 1 sec as we typically use [55] (*σ* = 1s). This flexible model offers better fitting of the HRF shape at the expense of more parameter estimations than the typical canonical hemodynamic response function [56]. Short-separation (SS) fNIRS signals are used as physiological regressors. The usual assumptions of the GLM are held. For each short-separation channel 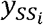, the GLM formulation in matrix format is as followed:

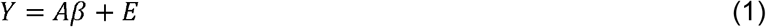

where 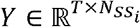 represents fNIRS recordings from all T time points and 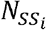 channels using 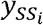 channel, 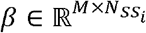 is the coefficient matrix with M regressors and 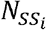 channels, 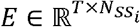 represents the residuals/noise term and *A* ∈ ℝ^*T×M*^ is the design matrix and is defined as followed:

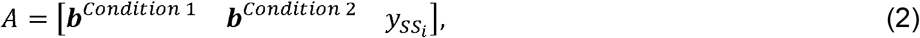

where **b**^*Condition i*^ ∈ ℝ^*T×H*^, *Condition i* ∈ {1,2} is a submatrix composed of the following column time vectors:

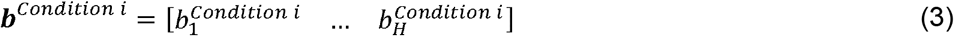

and each column vector 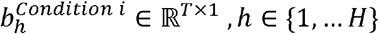 is defined as followed:

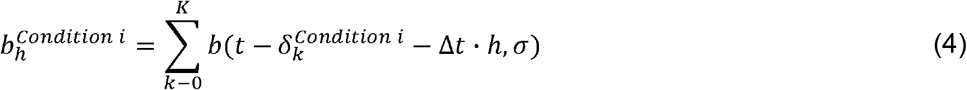

where K is the total number of trials in Condition i and 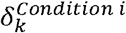 represents the timing of the onset of k^th^ stimulus. *b* (*μ, σ*) is the Gaussian function and is defined as followed:

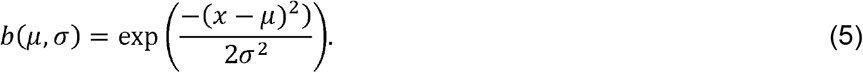

where both *μ* and *σ* are in seconds. Letting *A*^⊤^ be the transpose of *A*. The *β* coefficients in the GLM equation are solved using the ordinary least squares (OLS) method [57].

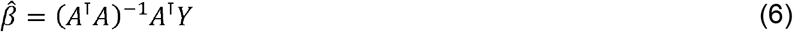

For the group-level HRFs, we average HRFs of all subjects for each channel and spatial location.

For the classification, fitting the GLM regression model to the entire data first before cross-validating the trials would result in information leakage [58]. In order to avoid leakage, we cross-validated both the GLM regression and classification steps. In each fold of the cross-validation, we fit the GLM model to the training dataset and estimated the coefficients using the OLS method. Then the SS coefficients 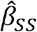 estimated from the training set are used for the test set where the individual trials are the difference between the measured fNIRS signals and the systemic physiological regressor weighted by the SS coefficients.

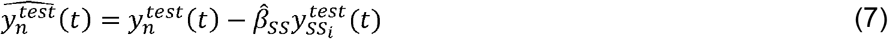

### Statistical Tests for GLM

For subject-level statistical test, to determine whether the hemodynamic activation significantly differed from 0, a one-tailed t-test was used on the estimated β-weights for the Gaussian temporal functions. We restricted ourselves to the β-weights with peaks between 1 and 8 seconds from the stimulus onset. The t-statistic was computed using the ratio of the sum of β-weight estimators to the square root of the variance of the sum of β-weights estimators. To determine whether the activations between two different conditions (e.g., left vs right) differed, the t-statistic was computed using the following contrast vector:

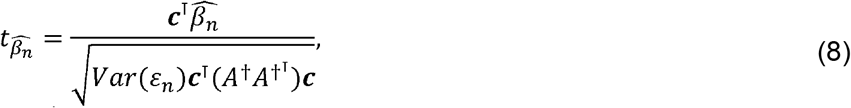

where *ε*_*n*_ is the error term for the n^th^ channel, ***c*** *∈* ℝ^*Mx*1^ is a contrast vector for two different conditions, *A*^†^ *∈* ℝ^*Mx*1^ is the Moore-Penrose inverse of the design matrix, 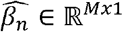 are the estimated β-weights for channel n. The Moore-Penrose inverse of the design matrix *A* assuming full column rank is:

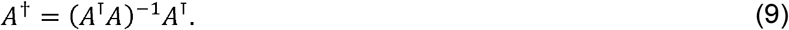

Let 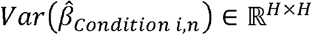 be the submatrix of 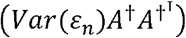 belonging to *Condition i*, and channel n. Then 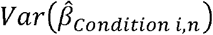 represents the covariance matrix for the HRF regressors for *Condition i*, and channel n. The 95% confidence intervals of the HRF for each condition and channel were computed as:

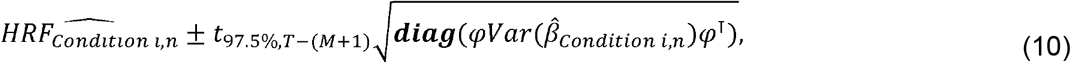

where we explicitly modeled independence between regressors by using only the diagonal elements of the matrix. 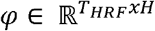 is the matrix of the standard Gaussian temporal basis functions with *T*_*HRF*_ as the time *φ* vector for the HRF. Then is constructed as followed:

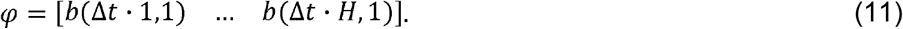

t_α,v_ is the t-value of a student’s t-distribution parameterized by the critical value α and the degree of freedom v.

For group-level statistical test, one-sample two-tailed t-test is used to determine whether the activation differed from 0 and two-sample two-tailed t-test is used to determine whether the activations between two conditions differed. Instead of using β-weights estimators, we defined the average of the HRF values over the range [1,8] seconds as a single observation for each subject, channel and condition. Then a t-test is performed over a sample of averaged HRF values for each channel. Independence between β-weight estimators is assumed.

To account for the large variation in the shapes of the HRFs, we provide an alternative group-level statistical test. We call this the aggregation method and will be described as followed. For each channel and condition, we count the percentage of subjects that are statistically significant at significance level α = 0.05, where equation (8) is used for statistical tests.

### fNIRS classification

In this study, to test the cross-validation (CV) accuracy of the classifier trained with the entire probe, we tested 2-class classification between left (−45°) and right (45°) spatial locations as well as 3-class classification between all 3 spatial locations. This classification will be termed all-channel classification. The features used are the area under curve of one-second-long segment of fNIRS signals. The classifier used is linear discriminant analysis with linear shrinkage of covariance matrix [59]. 5 other classifiers were also tested and their CV accuracies are reported in Supplementary Fig. S1. These are linear discriminant with non-linear estimator of covariance matrix using nuclear norm penalty [60], KNN [61] with the distance metric defined as 1 minus cosine similarity and 10 nearest neighbors, support vector machine [62-64] using cubic polynomial kernel, regularized with ridge penalty, random forest ensemble using 200 unpruned decision trees as weak learners, sampling 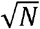 out of N channels for predictors and nonparametric bootstrapping [65], and gentle adaptive boosting using 200 unpruned decision trees as weak learners [66]. First, we tested [ΔhbO], [ΔHbR] and [ΔHbT] on all subjects and reported the grand average of CV accuracies. Next, we focused on [ΔHbT] and divided the subjects into two groups based on their task performance and reported their CV accuracies. The CV accuracies are reported for the interval 3-4 s after the cue onset in Fig. 5. We also plotted the correlations between subjects’ task performance and individual subjects’ CV accuracies using [ΔHbT] over interval 3-4 s after the cue onset (Fig. 6)

**Figure 4.**
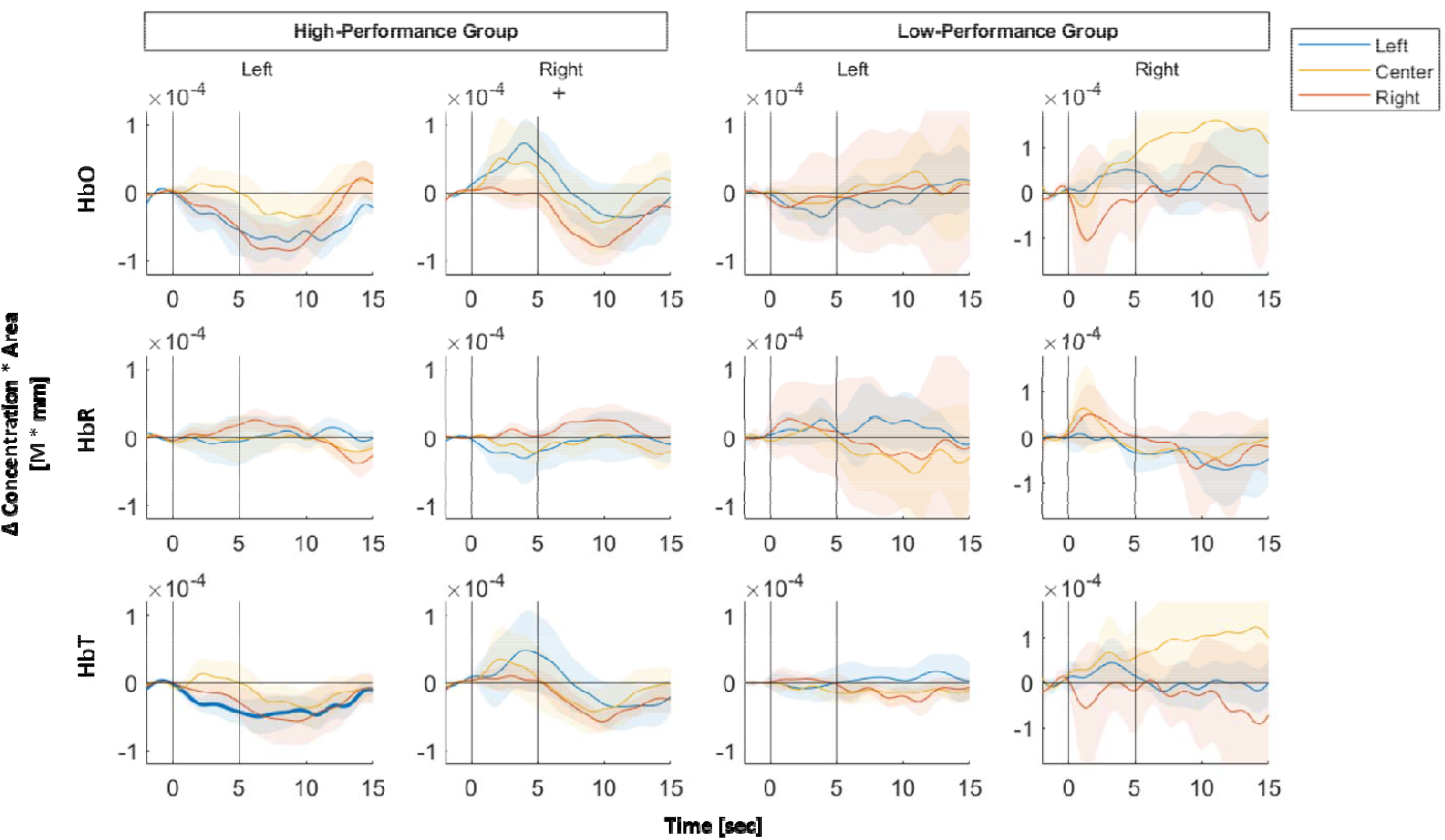
HRFs of FEF by task performance. Same as Figure 3 but for group average of a high and a low-performing groups. Range in M*mm of y-axis is [-1.2e-4 1.2e-4] for the 3 left columns but [-1.8e-4 1.8e-4] for the rightmost column.

**Figure 5.**
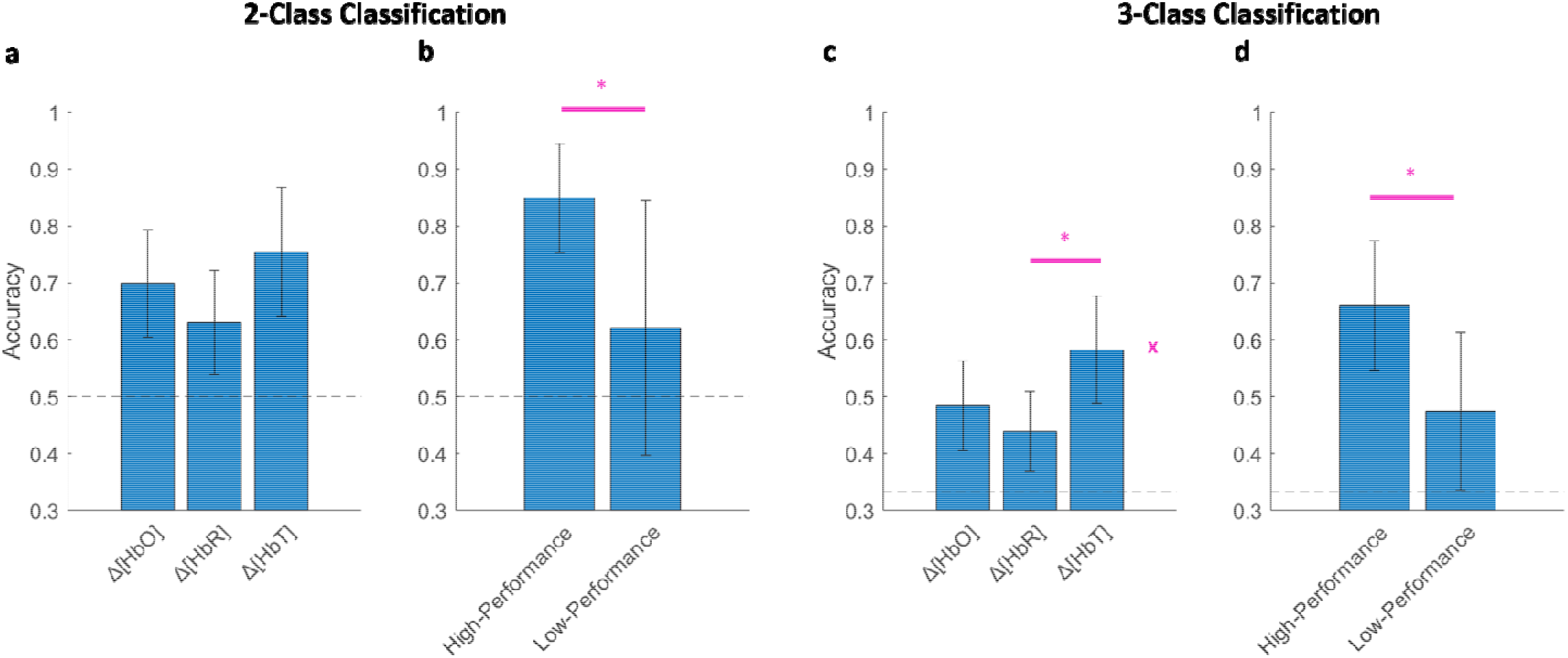
Δ[HbT] has the highest CV accuracies, CV accuracies are highly dependent on task performance. (a) 2-class grand-average CV results for all 12 subjects at t = 3-4s. Error bar represents 95% confidence interval. x represents F-test using one-way ANOVA with significance of 0.05. * represents significance level of p < 0.05. (F_2,33_ = 1.86, p = 0.17). (b) Focusing on Δ[HbT], 2-class CV accuracies shown for high-performance and low-performance groups. Two-sample two-tailed t-test: t(10) = 2.79, p = 0.019. Similar plots are shown for 3-class classification in (c) and (d). (c) One-way ANOVA F-test, F_2,33_ = 3.87, p = 0.031. (d) Two-sample two-tailed t-test: t(10) = 2.69, p = 0.023.

**Figure 6.**
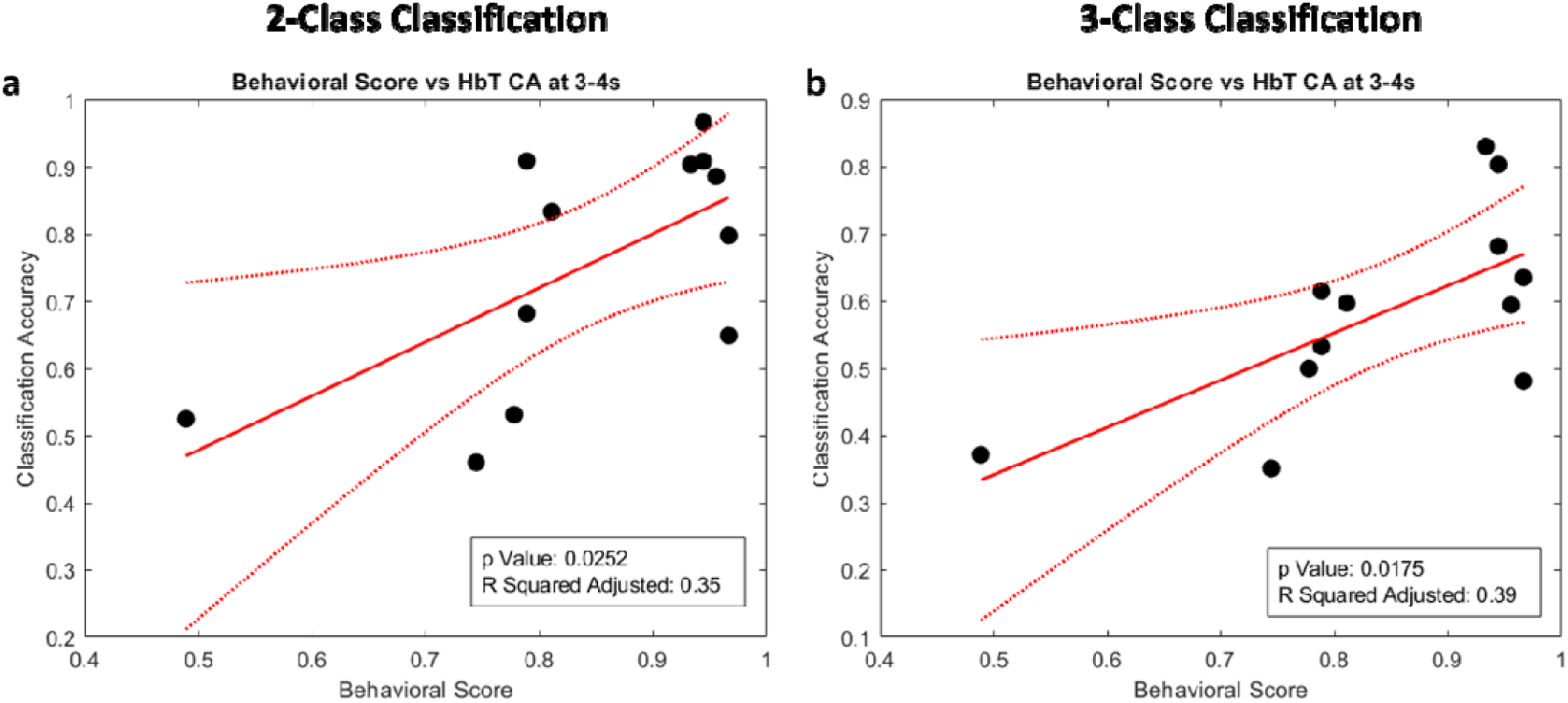
Strong correlation between CV accuracies and task performance. (a) Linear regression between behavioral score and 2-class CV accuracy for Δ[HbT] using window 3-4s. (Slope coefficient: t(10) = 2.63, p=0.025, adjusted r^2^ =0.35) (b) Linear regression between behavioral score and 3-class CV accuracy for Δ[HbT] using interval 3-4s. (Slope coefficient: t(10) = 4.84, p=0.018, adjusted r^2^ =0.39). In all cases, n = 12. Dashed lines represent 95% confidence intervals.

To assess the tradeoff between the number of channels, latency, and cross-validation accuracies, we plotted the CV accuracies of classifiers trained with different regions of the probe as a function of different window length in Fig. 8. All windows tested started at cue onset. To account for the variation in reported MNI locations of different ROIs, as well as the variation in head and brain sizes and ROIs’ actual locations, neighboring channels are aggregated to form 5 different subsets. For example, a subset named left FEF (L FEF) has 10 channels and covers FEF and the following nearby ROIs: iPCS, dlPFC, tgPCS, and sPCS in the left hemisphere. Similarly, right FEF (R FEF) has 10 channels and covers FEF and nearby ROIs in the right hemisphere, left STG subset has 1 channel and covers left STG region, right STG subset has one channel and covers right STG region, left IPS subset has 4 channels and covers IPS2-4, mSPL, antIPS and SPL1 in the left hemisphere and similarly for right IPS (Fig. 2).

To quantify the contribution of each channel, we excluded it from the all-channel classification and then used it to train a classifier and retrieve its CV accuracy for window t=3-4s. This will be called leave-one-feature-out (LOFO) classification. Specifically, we took the difference in CV accuracies between all-channel classification and LOFO classification and defined it as the LOFO impact [67]. Finally, we ran single-channel classification for windows t=3-4s where a single channel is used to train a classifier (Fig. 7). This is done for all channels. We used 10 repetitions of 5-fold cross validation in all cases except for linear discriminant with nonlinear estimator of covariance matrix, which used 10 repetitions of 5-fold nested cross validation. In all cases, 1-0 loss function was used. For the confidence interval of the means of cross-validation accuracy, we used a confidence level of 95% using student’s t-distribution.

**Figure 7.**
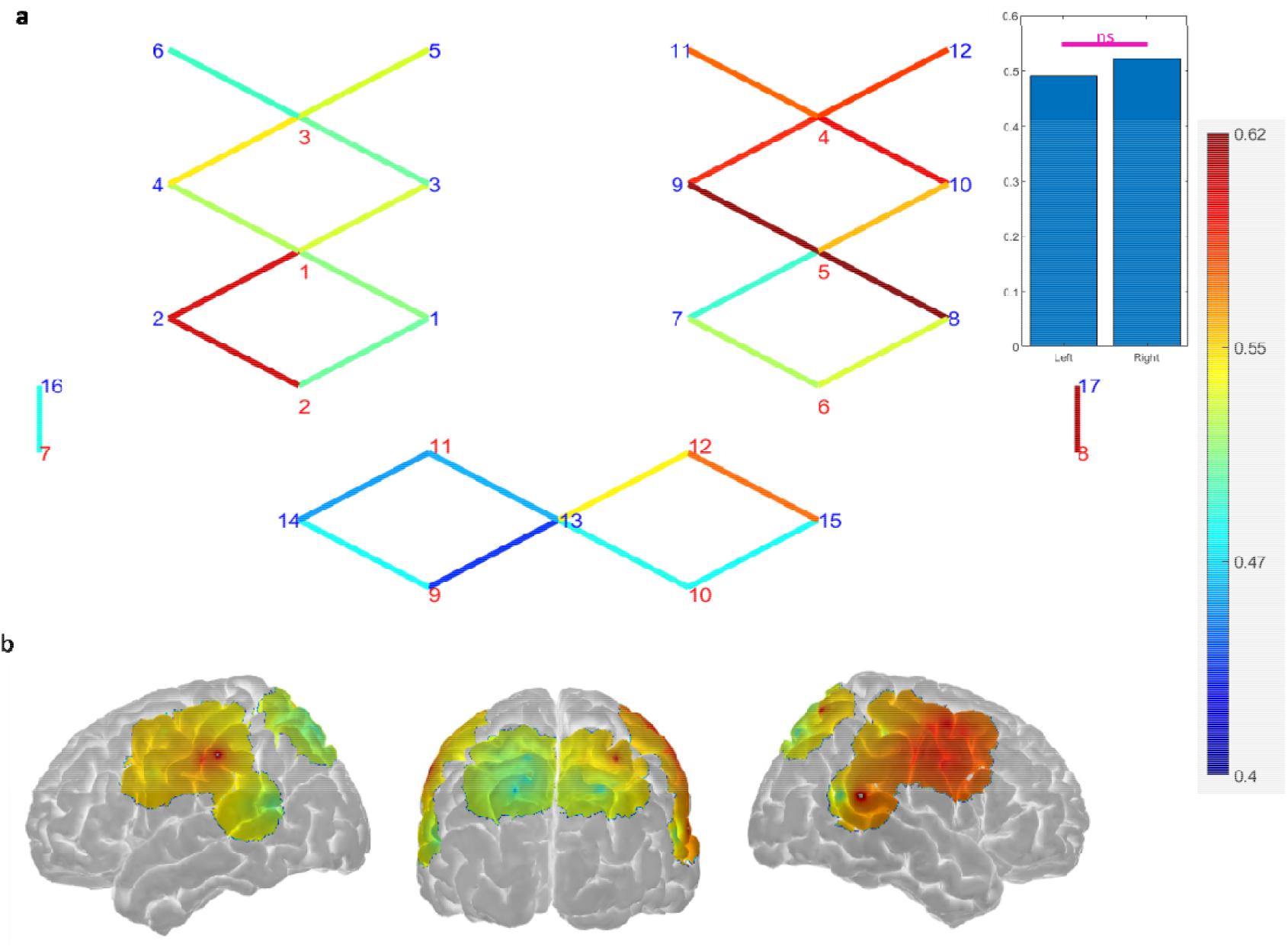
Single-Channel CV Accuracies. (a) 2-D probe layout where each line represents a channel with the corresponding colormap indicating their CV accuracies for window t=3-4s (during movie presentation). Red numbers represent source optodes and blue numbers represent detector optodes. Inset: single chancel classification grouped by hemisphere, reported as mean and standard deviation. Left: 0.51±0.047. Right: 0.56 ±0.050. ns is not significant. Two-tailed two-sample t-test for lateralization: t(28) = -1.81, p=0.081. (b) Colormap of the single-channel CV accuracies plotted on the surface of the brain with interpolation. Mean and median of 95% CI across all channels are 0.168 and 0.135 respectively.

### Statistical Tests for Classification

To test for statistical significance between 3 different chromophores, we used one-way ANOVA test. If the overall F test passed the given significance of 0.05, we followed up with post-hoc two-tailed two-sampled t-tests and used Bonferroni-corrected p-values. To test for statistical significance in decoding performance between high and low-performance group, we used two-tailed two-sampled t-tests.

## Results

### Behavioral Results

All 12 subjects performed significantly above chance level in the task (mean ± standard deviation: 84.3±14.1%). Of these 7 subjects, who had performance levels of 80% or above were categorized as high-performing, and 5 subjects with performance levels less than 80% were categorized as low performing.

### HRFs

We plot the HRFs for various conditions, spatial locations and chromophores in the figures below. One example channel that shows different HRF patterns for left and right spatial locations is located in the FEF region in the right hemisphere and is shown in the right column of Figure 3. In addition, this channel has the highest correlation coefficient between task performance and single-channel CV accuracies. Its contralateral channel (in the left hemisphere) is also shown in the left column of Figure 3. In addition, we observed mostly a positive hemodynamic response in the right hemisphere and mostly a negative hemodynamic response in the left hemisphere in Δ[HbO] and Δ[HbT]. The different magnitudes of the responses to spatial location suggests that it’s possible to decode the spatial location using features of the HRF. In addition, we tested for statistical significance and only found that the Δ[HbO] activation in the right FEF in response to the left spatial location was statistically significant at α = 0.05 (t(11)=2.73, p = 0.0196).

To investigate if there were any differences between the high-performance and low-performance groups, we examined the group-averaged HRFs. Figure 4 shows example group averaged HRFs from the FEF region. The most striking pattern was that the response variabilities (as shown with 95% confidence intervals) in the HRFs are substantially wider in the low-performance group compared to the high-performance group. Moreover, the positive hemodynamic response in the right hemisphere and negative hemodynamic response in the left hemisphere was more pronounced in the high-performance group. Statistical tests showed that Δ[HbT] activation on the left FEF in response to the left spatial location in the high-performance group is statistically significant at α = 0.05, (t(6) = -2.64, p = 0.039). In addition, the difference in Δ[HbO] activation between left and right spatial location in right FEF channel for high-performance group is statistically significant.

Due to large subject variability and the small sample size, Supplementary Table 1 provides the aggregation method where we calculated the percentage of subjects that achieved subject-level statistical significance using α = 0.05. In all but one of the cases, subjects from the high-performance group are more likely to achieve statistical significance than subjects from the low-performing group.

### Classification/Decoding Performance

Next, we plotted the decoding performance using all-channel cross validation (CV) accuracies. Figure 5a and c show the averages of 2-class and 3-class CV accuracies, respectively, for the interval t=3-4s after the cue onset when using all channels in the classification. In both cases, CV accuracies are significantly above chance level and are highest when using Δ[HbT] and lowest when using Δ[HbR]. Next, we focused on Δ[HbT] and divided the subjects into high-performance and low-performance groups. We found that decoding performance was significantly higher in high performance subjects compared to low performance subjects (Fig. 5b and d). To further quantify the relationship between CV accuracies and task performance, we plotted these quantities in Fig. 6a-b and performed linear regression when classifying using Δ[HbT] between 3 and 4 seconds. The significant correlation between the CV accuracies and task performance indicates that the decoding performance is dependent on the subject’s task performance.

We next focused on CV accuracy in the high-performance group using Δ[HbT] signals. To assess the contribution of specific brain regions towards decoding performance we quantified single channel classification performance. Fig. 7 shows the single-channel 2-class classification using a 2D channel layout (a) and interpolated over the brain surface (b) for the interval 3-4s after the cue onset. On average, decoding performance was higher in the right hemisphere but the difference was not significant at the 0.05 level (Fig. 7a inset). In addition, the top 2 performing channels are located in the right FEF region. The CV accuracy from single-channel classification (Fig. 7) was lower compared to all-channel classification (Fig. 5b).

In order to examine the contribution of each subset of channels to the CV accuracy as well as the tradeoff between the number of channels and window length, we plotted 2-class CV accuracies for left and right FEF and IPS regions (Fig. 8). All-channel classification is shown for comparison. All windows start at the cue onset. For the all-channel classification, the cross-validation accuracy is already significantly above chance level using 1 sec window length (t(6) = 2.94, p = 0.026). In addition, the lateralization of decoding performance to the right hemisphere is evident in both FEF and IPS regions. However, FEF has substantially higher decoding performance than IPS. Even when combining left and right IPS regions for a total of 8 channels to account for FEF’s higher dimension, it remains at chance level (not shown). Seeing that the R FEF region is the primary driver of decoding performance, in order to identify specific key channels within the R FEF region, we performed LOFO-classification for all channels (Supplementary Fig. S2). All top 3 channels within the R FEF region are located closer to the medial axis (Inset of Fig. 8). Specifically, these channels cover FEF and superior precentral sulcus. While FEF and IPS both are part of the dorsal frontoparietal network known to play an important role in visuospatial attention, our study shows that FEF has substantially better fNIRS decoding performance than IPS.

**Figure 8.**
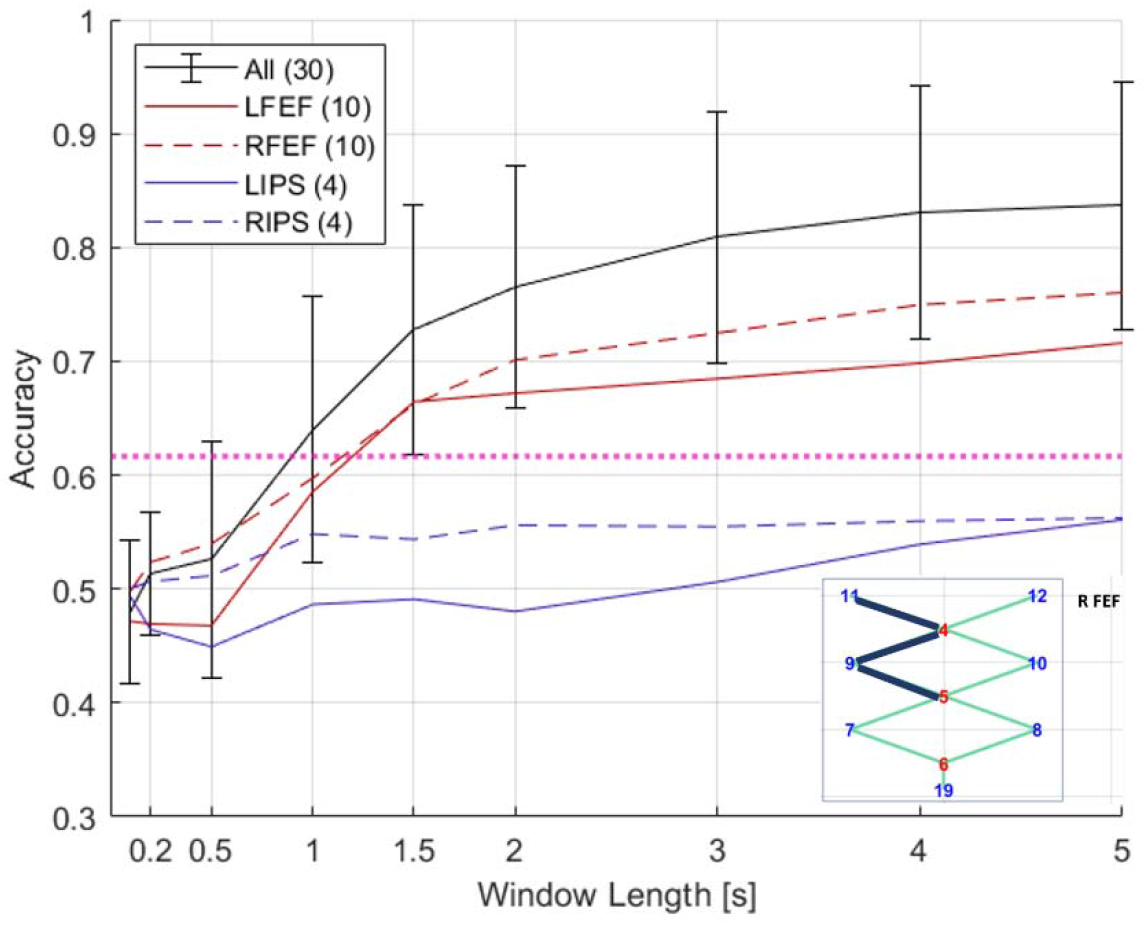
Time course of improvement in CV accuracy. Average classification accuracies of high-performing group as a function of time. All windows start at 0s (cue onset). Decision window lengths tested here are 0.1s, 0.2s, 0.5s, 1s, 1.5s, 2s, 3s, 4s, and 5s. LFEF = left frontal eye field. RFEF = right frontal eye field. LIPS = left intraparietal sulcus. RIPS = right intraparietal sulcus. The numbers next to subregion in legend represent the number of channels. Pink line indicates statistical significance for p = 0.05 using two-tailed t-test and the given standard deviation from our cross validation accuracies across subjects at window length 1s. Error bars represent 95% confidence intervals. Error bars for other lines are similar in magnitude and are not shown for visual clarity. (Inset) Top 3 channels from R FEF based on LOFO impact are shown as navy lines.

## Discussion

In this study, we demonstrated the capability of fNIRS to decode attended spatial location in CSA. During quantitative assessment of the classifier, we made several discoveries. First, we found that Δ[HbT] provides substantially higher CV accuracy than Δ[HbO] and Δ[HbR], consistent with a previous report [68]. Second, the CV accuracy of a classifier was significantly correlated with behavioral performance. It is well known that attention can modulate cortical response properties, e.g., in fMRI studies on auditory primary and secondary cortices [69,70]. The difference in CV accuracy between high and low performance groups supports the idea that attention improves decoding performance. Third, we identified right FEF region as making the largest contribution to overall decoding performance. Interestingly, FEF had substantially higher decoding performance compared to IPS even though both are part of dorsal frontoparietal network, which has been implicated in previous studies of spatial attention [30-32]. Finally, we found that CV accuracy improved faster than expected, reaching significantly (at α = 0.05) above chance level at 1s, well before of the peak of the HRFs. This suggests that the onset of the HRFs, not just the peaks which occur much later, can contain useful information for classification. A contributing factor in the increased classification performance in high performing subjects is the lower variability of the underlying HRFs in high performing subjects (Figure 4). Signal detection theory predicts that higher detection and discrimination performance can result from an increase in the mean difference and/or a decrease in the variability of the two distributions [71]. A previous study found reduced variability of neural responses in auditory cortex associated with improved neural detection during auditory task performance [72]. Such an effect is a potential neural mechanism underlying the HRFs in our experiments.

One advantage of fNIRS over EEG is better spatial resolution, which revealed FEF to be a critical region for decoding the attended location in CSA. Moreover, relative to EEG, fNIRS is less influenced by eye movement and blinking, enabling the use of visual stimuli for decoding brain signals without significant contributions from eye movements. In the future, it would be interesting to use an integrated fNIRS-EEG system to leverage the high spatial resolution of fNIRS and the high temporal resolution of EEG to develop a robust and rapid decoding algorithm. It also would be interesting to see whether fNIRS can be applied in assistive devices to help those struggling with CSA, e.g., populations with ADHD, autism and hearing impairment.

## Data Availability

Code and data are available at https://github.com/NoPenguinsLand/fNIRSCodes_Manuscript.

## Acknowledgements

MN want to express his gratitude to Parya Farzam, Aneka Pradhan, Vibhav Jha and Ahona Dev for their assistance with experimental setup and data collection.

## Author contributions

AL: methodology and writing – review and editing. DB: methodology and writing – review and editing. KS: conceptualization, project supervision and writing – review and editing. MN: methodology, experimental setup, data collection, algorithm and data analysis, figures generation and data visualization, writing – original draft, writing – review and editing. MY: methodology, project supervision and writing – review and editing.

## Competing Interests

The authors declare no competing interests.

Correspondence and requests for materials should be addressed to KS.

