## Supplementary Figures and Table for "Decoding Attended Spatial Location during Complex Scene Analysis with fNIRS"

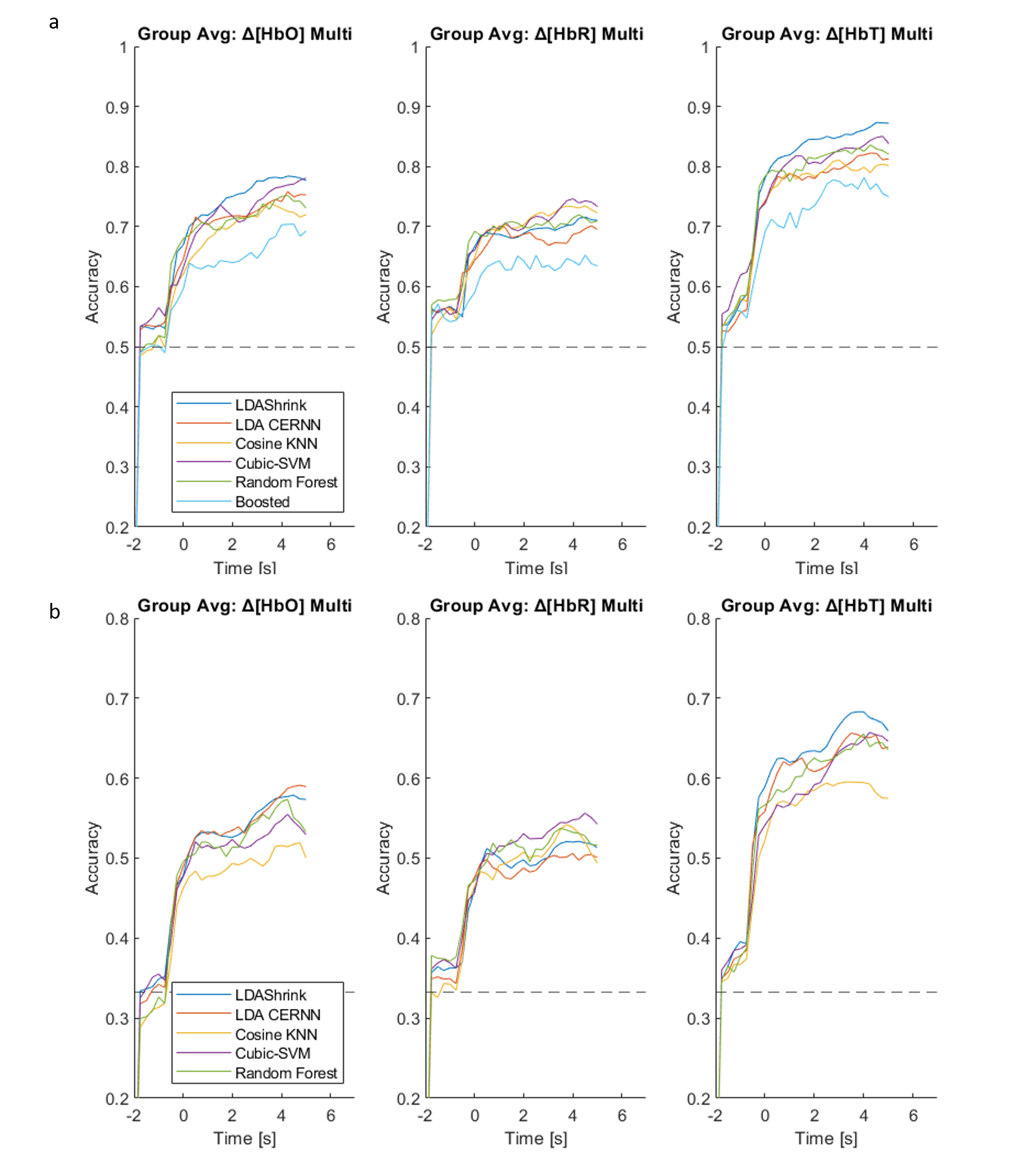

Supplemental Figure 1 (a) 2-Class CV accuracies for Δ[HbO] on the left panel, Δ[HbR] on the middle panel, and Δ[HbT] on the right panel. Error bars representing 95% confidence interval are omitted for clarity. (b) Same as (a) but for 3-class CV accuracies. It’s very noteworthy that linear discriminant analysis with linear shrinkage has the highest classification accuracy for most of the trial segment for Δ[HbT].

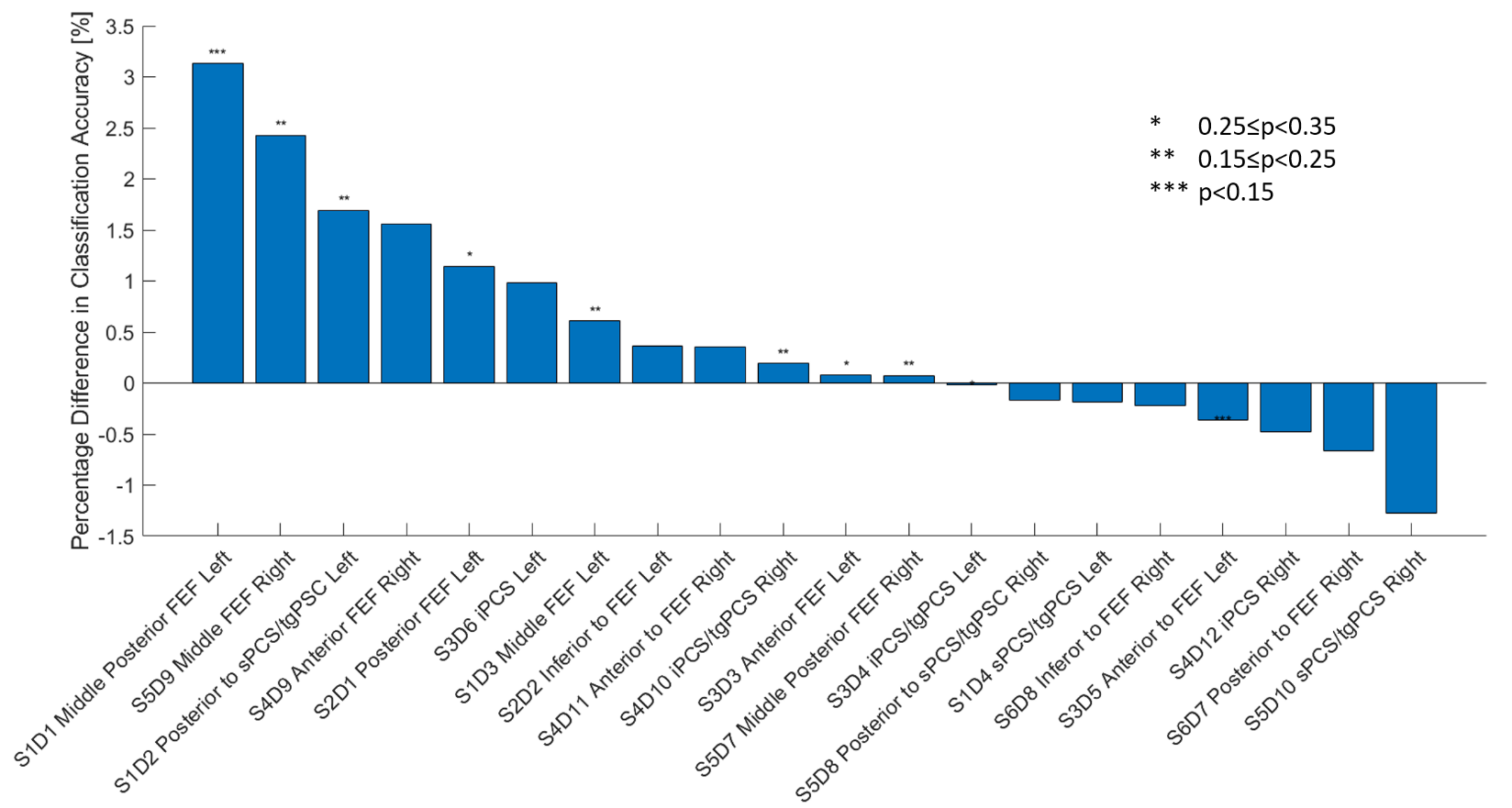

Supplemental Figure 2. FEF has biggest impact as determined by leave-one-feature-out classification. Bar chart showing the averaged percentage difference in CV accuracies between all-channel classification and leave-one-channel-out classification.

| (High-Performance Group, n=7) | | | | | | | | |
| --- | --- | --- | --- | --- | --- | --- | --- | --- |
| HbT | Left | Right | Center |  | HbT | L vs R | L vs C | R vs C |
| L FEF | 85.71 | 85.71 | 71.43 |  | L FEF | 85.71 | 71.43 | 57.14 |
| R FEF | 100.00 | 71.43 | 85.71 |  | R FEF | 71.43 | 85.71 | 100.00 |
| (Low-Performance Group, n=5) | | | | | | | | |
| HbT | Left | Right | Center |  | HbT | L vs R | L vs C | R vs C |
| L FEF | 60.00 | 40.00 | 40.00 |  | L FEF | 40.00 | 60.00 | 40.00 |
| R FEF | 80.00 | 40.00 | 60.00 |  | R FEF | 80.00 | 20.00 | 60.00 |

Supplementary Table 1. Aggregation Method. Percentage of subjects with statistical significance for both activation (using one-sample one-tailed t-test) and difference in activation between two spatial locations (using two-sample one-tailed t-test). High-performance group is more likely to yield statistical significance in all except one case. All use significance of 0.05.
